# Enrichment of halogenated organic compounds and their degrading microorganisms in the deepest ocean

**DOI:** 10.1101/2025.02.11.637578

**Authors:** Rulong Liu, Hui Wei, Zhiao Xu, Yuheng Liu, Zhixuan Wang, Jiani He, Min Luo, Federico Baltar, Yunping Xu, Qirui Liang, Liting Huang, Li Wang, Jiasong Fang

**Affiliations:** College of Oceanography and Ecological Science, Shanghai Ocean University, Shanghai, China; Department of Functional and Evolutionary Ecology, University of Vienna, Vienna, Austria; Laboratory for Marine Biology and Biotechnology, Qingdao National Laboratory for Marine Science and Technology, Qingdao, China; Department of Natural Sciences, Hawaii Pacific University, Honolulu, HI, USA

**Author notes:** Correspondence: Rulong Liu, Li Wang, Jiasong Fang. these authors contribute equally (Hui Wei, Zhiao Xu).

## Abstract

The hadal trenches, the deepest regions of the ocean, serve as the final resting place for marine particles and as “tunnels” for material exchange between the ocean and Earth’s interior. Despite their extreme conditions, the trenches contain high content of organic carbon (OC) and active microbial carbon turnover, are hotspots for deep-sea OC degradation and unique microbial processes. However, little is known about the OC components and microbial metabolisms driving the OC degradation in the trenches. This study reveals unexpectedly high concentrations of halogenated organic compounds (HOCs) in the Mariana Trench sediments, with about 1 halogen per 58 carbon atoms in the OC pool. Systematic analysis of deep-sequenced metagenomes and metagenome-assembled-genomes from the global ocean shows significantly higher abundance of the genes for biodegradation of HOCs (dehalogenation) in trench microbes, with microorganisms capable of dehalogenation belonging to 16 phyla and 52 orders, 75% of which identified as HOCs degraders for the first time. Microcosms simulating trench conditions showed rapid degradation of typical HOCs and transcription of genes related with HOCs metabolisms, demonstrating the active HOCs degradation by the trench microorganisms. The findings suggest the HOCs metabolism as an important process in the OC remineralization of deep-sea trenches, advancing understanding of deep-sea carbon cycling and microbial survival strategies.

## Introduction

The deep ocean, covering 65% of the Earth’s surface and housing 75% of the ocean’s prokaryotic biomass, plays a crucial role in the remineralization and long-term storage of marine organic carbon ^1^. However, our understanding of the metabolism of deep-sea microorganisms and the carbon turnover processes they drive remains limited, especially concerning carbon cycling in the ocean’s deepest regions, known as hadal trenches. The hadal trenches, which are found at depths exceeding 6,000 meters, are among the most inaccessible and least understood marine environments on the planet ^2, 3^. Located in subduction zones, the hadal trenches serve as both the endpoint for sinking organic matter from the upper ocean and as a “channel” for material exchange between the ocean and the Earth’s deep interior ^3^. Recent studies have shown that the hadal zone is rich in organic matter and exhibits unexpectedly active microbial organic carbon turnover, emerging as “hotspots” for organic carbon remineralization in the deep ocean ^2, 4^. Furthermore, the extreme conditions of the hadal zone, such as high pressure and geographic isolation, have led to the development of unique microbial communities and special metabolic processes, forming a distinct “trench biosphere” ^5, 6^. Understanding the processes and mechanisms by which hadal microbes drive the degradation of sedimentary organic carbon is essential for advancing knowledge of carbon cycling and life processes in deep-sea environments ^2,3^.

While studies have demonstrated that hadal trench microorganisms exhibit potential for chemoautotrophic metabolism such as ammonia oxidation ^7^, metagenomic and genomic analyses suggest that microbial taxa present in the trench sediment predominantly adopt a heterotrophic lifestyle, possessing metabolic genes and pathways for degradation of a wide range of organic matter, including simple sugars, peptides, fatty acids, proteins and other organic detritus ^6, 8^. Particularly, most of the studies demonstrated the metabolic potential of hadal trench microbes to utilize multiple complex and recalcitrant organic compounds, such as hydrocarbons, polysaccharides and aromatic compounds ^6, 8^. These findings are consistent with the higher humification of organic carbon in the bottom of hadal sediments ^9^, suggesting that the utilization of recalcitrant organic carbon might be a crucial metabolic strategy for hadal microbes ^8, 10^. However, only a limited number of these metabolic processes, such as hydrocarbon degradation, have been experimentally validated combined evidences of biogeochemical detection of substrates and the assessment of microbial degradation activities ^11, 12^. Due to the diverse and variable sources of organic matter^3^, along with the unique biological, geological, and physicochemical factors regulating the dynamics of materials in hadal trenches ^13, 14^, the composition of organic matter in hadal sediments is highly complex. Consequently, our understanding of the components of organic carbon (OC) in hadal trenches and the microbial processes driving OC degradation in these sediments remains limited.

Halogenated organic compounds (HOCs) are widely recognized as environmentally hazardous substances, raising significant global concerns. As persistent organic pollutants, most anthropogenic HOCs exhibit high toxicity to various organisms and a strong tendency to bioaccumulate, causing considerable harm to the marine ecosystems ^15^. In addition to human sources, large quantities of natural HOCs are also produced through various biological and abiotic processes ^16, 17^. The ocean, being rich in halogen atoms, serves as the largest source of natural HOCs on Earth ^18^, with high level of HOCs detected in sinking particles ^19^. Due to their recalcitrant nature, HOCs are thought to be transported to the deep ocean via mechanisms such as biological pump and ultimately sequestered in deep-sea sediment ^15, 19, 20^. However, little is known about the content of HOCs and their fate in the deep ocean.

Hadal trenches serve as major depositories of different types of pollutants ^21, 22^. Recent studies have identified a variety of HOCs, such as organochlorine pesticides dichlorodiphenyltrichloroethane (DDT) and chlordane, industrial pollutants PCB and polybrominated diphenyl ethers (PBDE), in the macro-organisms and sediments of the Mariana Trench ^22, 23, 24, 25^. In addition, our previous work revealed that genes encoding complete metabolic pathways for HOCs degradation are present in genomes of hadal Chloroflexi, one of the dominant taxa in in trench sediments ^10, 26^. The finding suggests the possibility of microbial HOCs degradation in the hadal trench sediment. Nevertheless, the metabolic processes, degradation activities as well as the significance of the HOCs degradation in hadal trench carbon cycling is not explored. In this study, we applied an optimized combustion-IC based approach to determine the content and composition of total HOCs in sediments of the Mariana Trench. Surprisingly, the results revealed that the total HOCs represent a large proportion of OC pool in the trench sediments. Systematic metagenomic and genomic analysis further showed that genes encoding enzymes and pathways for HOCs degradation are enriched and prevalent in hadal trenches sediments, and are widely distributed across more than half of the microbial taxa. Microcosm experiments simulating trench conditions further demonstrated active degradation of HOCs by trench microbial communities. These findings provide novel insights into the significance of HOCs and their microbial degradation processes in carbon cycling of the hadal zone.

## Results and Discussion

### High content of HOCs in hadal trench sediments

A method based on combustion-ion chromoatography were developed and optimized to show high accuracy, precision and reproducibility in determining the content of adsorbable organic halogens (AOX) including fluorine, chlorine and bromine (AOF, AOCl, AOBr), and the corresponding insoluble organic halogens (IOF, IOCl, IOBr, IOX), as well as total organic halogen contents (TOBr, TOCl, TOF, TOX) in marine sediments ^27^. In this study, the content and composition of HOCs were determined in 38 sediment samples of 4 sediment cores taken from 5842-10901 meters of the Mariana Trench, which resulted in a total number of 430 data points of different components of HOCs (Fig. 1; Fig. S1; Table S1). The results showed that HOCs presented in all of the trench sediments tested (Fig. 1B). The values of AOX, which indicates dissolved HOCs adsorbable by activated carbon, ranged from 4.90±0.70 to 383.14±64.17 mg/kg, while the insoluble HOCs (IOX) ranged from 71.82±0.50 to 459.96±21.90 mg/kg (Fig. 1B), were significantly higher than the content of AOX (p < 0.01, t-text). The dissolved fraction of HOCs were mainly dominated by organic chlorine compounds (AOCl), which were significantly more abundant than AOBr and AOF. In contrast, organic fluorine compounds (IOF) showed the highest abundance in the insoluble HOCs, followed by IOCl (Fig. 1B). Organic bromine compounds showed the lowest abundance in both the dissolved and insoluble HOCs (Fig. 1B). The concentration of total organic halogen (TOX) in sediments of the Mariana Trench ranged from 110.33 ± 2.71 to 594.15 ± 15.28 mg/kg, with an average value of 292.47 mg/kg (Fig. 1B). Although hydrocarbons and lipid biomarkers, e.g. alkanes, carboxylic acids, alcohols and GDGTs, have been reported to be potential components in OCs of hadal trench sediments, these compounds generally showed low abundance (up to 15 mg/kg) ^11, 12, 28^. HOCs determined in this study were at least one magnitude higher than the reported OC compounds, making HOCs the largest known component of organic matter in hadal trench sediments. The contents of total organic carbon were relatively stable in the tested samples, ranged from 0.48 – 0.69% (0.57 ± 0.06 %) (Fig. 1B). As a result, the calculated molar ratios of carbon to halogen atoms (C:X) were ranged from 17.8 to 121.8, with an average value of 49.2. This indicates an average of one halogen atom for every 49.2 organic carbon atoms, suggesting a high proportion of HOCs within the organic carbon pool in the tested trench sediments.

**Figure 1.**
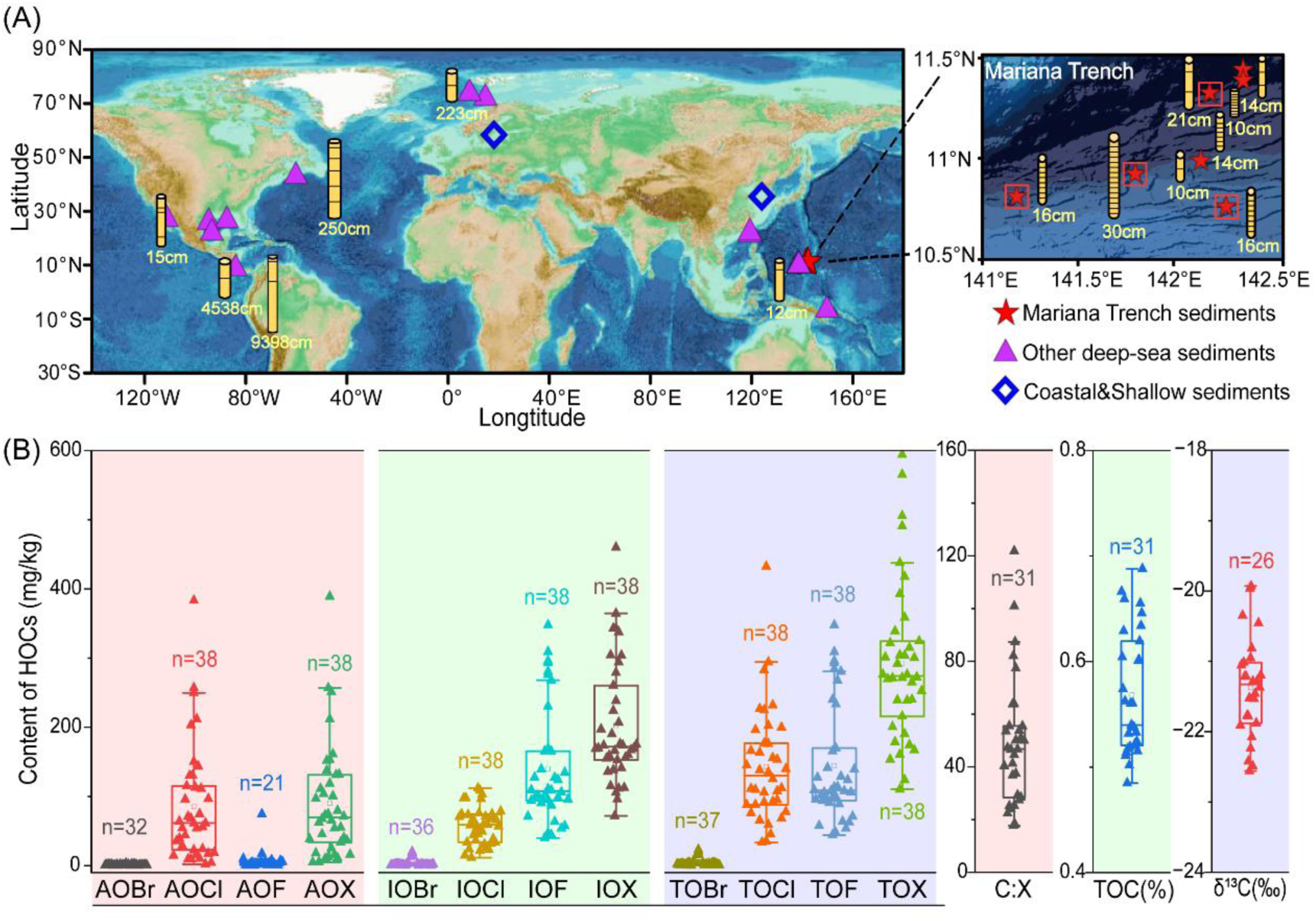
(A) Locations of sediment cores involved in the analysis of this study. The red stars with squares indicate the stations and samples with detailed geochemical data analyzed. (B) The geochemical parameters of the sediment cores from the Mariana Trench, including the concentrations of adsorbable organic halogens (AOF, AOCl, AOBr and AOX), and the corresponding insoluble organic halogens (IOF, IOCl, IOBr, IOX and IOX), as well as total organic halogen contents (TOBr, TOCl, TOF, and TOX). The values of total organic carbon (TOC) and their carbon isotopic compositions (δ13C), as well the molar ratio of calculated molar ratios of organic carbon to halogen atoms (C:X) were also shown.

Currently, understanding on components of deep-sea organic matter is scarce, and to the best of our knowledge, only one existing study reported the TOX values in the deep ocean and from only two stations ^27^. The values of TOX in sediments of the MT were significantly higher than those reported in coastal and other deep-sea regions, despite the lower TOC content in the trench sediments (Fig. S2) ^27^. Consequently, the proportion of HOCs were significantly higher in organic carbon of the trench sediments than those in the coastal or other deep-sea sediments (evidenced by the lower C:X ratio) (Fig. S2).

HOCs in marine environments might be natural or anthropogenic sourced ^17, 18^. The hadal trenches have been reported to accumulate many types of manmade pollutants such as black carbon ^29^, mercury ^30^, and plastics ^21^. It was therefore not surprising to have the anthropogenic HOCs in the trenches, and indeed, various types of anthropogenic HOCs, such as PCBs, PBDEs, and halogenated organic pesticides have been reported in animals of several trenches, including the MT ^22, 23, 24, 31^. However, the reported anthropogenic HOCs generally showed low concentration in the trench sediment, with average values ranged from 0 to 2 µg/kg ^22, 23, 24^. Our results showed that the concentration of TOX were magnitudes higher than all of the reported anthropogenic HOCs, suggesting that large amount of HOCs might be come from other sources. Carbon isotopic compositions (δ13C) of TOC ranged between -20.0 to - 22.6 ‰, with an average value of -21.4 ‰ in the sediments analyzed (Fig. 1B), which indicating that the organic matter in the sediments of the MT was mainly marine sourced ^32^. Such finding was consistent with conclusions driven by previous studies carried out in the same region using the carbon isotopic or biomarker evidences ^32, 33^. Marine sourced HOCs might therefore be an important contributor for the measured high content of total HOCs in the hadal sediment of the MT. In fact, previous studies have shown that HOCs can be formed biologically via synthesis of marine organisms or abiotically via physical/chemical halogenation of natural organic matter in seawater^17, 18, 19^, and that the marine particles can be a natural sink for HOCs ^19^. The TOX values determined in the trench sediments here were comparable with those in deep-sea particles (180-694 mg/kg) ^19^, supporting the potentials of contribution from marine particles.

### Enrichment of dehalogenase in sediments of the Mariana Trench

HOCs are typically considered to be resistant to microbial degradation ^16^. The potential for microorganisms residing in hadal trench sediments to metabolize and degrade HOCs was investigated through a comprehensive metagenomic analysis focusing on the occurrence, relative abundance, and distribution of dehalogenase, which is the key enzyme for the biodegradation of HOCs ^34, 35, 36^. A total of 54 metagenomes from global ocean sediments were examined, comprising 27 deeply sequenced metagenomes from sediment cores collected from depths of 5,437 to 10,954 meters in the Mariana Trench, alongside 27 metagenomes from other deep-sea and shallow coastal sediments (Fig. 1A, Table S2). Following sequence preprocessing and functional annotation against the KEGG database, various types of dehalogenase genes were identified and validated through BLASTP search against the UniProt protein database. The relative abundance of these confirmed dehalogenase genes was quantified and expressed as genes per million (GPM). The results indicated the presence of dehalogenases in all metagenomes derived from Mariana Trench sediments (Fig. 2A). Specifically, four distinct types of dehalogenases were identified: haloalkane dehalogenase, S-2-haloacid dehalogenase, haloacetate dehalogenase, and reductive dehalogenase (Fig. 2A). These enzymes facilitate the biodegradation of various HOCs through either hydrolytic or reductive mechanisms ^18^, suggesting a significant metabolic versatility regarding HOCs within trench sediments. Notably, the relative abundance of haloalkane dehalogenase and S-2-haloacid dehalogenase was significantly higher in Mariana Trench sediments compared to other deep-sea and shallow sediments (p < 0.001, t-test) (Fig. 2A), indicating an enrichment of dehalogenation metabolism within the microbiome of the Mariana Trench. These results align with the high levels of HOCs found in the sediment of the trench (Fig. 1B), implying that trench microorganisms may utilize HOCs as substrates for living. Furthermore, phylogenetic analysis of 2,300 haloalkane dehalogenase sequences from the Mariana Trench metagenomes, in conjunction with all manually annotated and non-redundant haloalkane dehalogenase sequences available in the UniProt-Swiss database, revealed that the microbial communities in the Mariana Trench encompass all three types of haloalkane dehalogenases (HDL-I, II, and III) (Fig. 2B). This suggests functional complementation among these enzymes, as different types of the enzyme exhibit varying substrate specificities ^37^. Interestingly, the sequences derived from the trench metagenomes formed unique clusters that were distinct from known haloalkane dehalogenases (Fig. 2B), indicating the presence of novel phylotypes that may possess unique structural and functional characteristics, which were potentially resulting from long-term adaptation to the extreme environmental conditions of the hadal trench, such as low temperatures and high pressures ^2, 3^.

**Figure 2.**
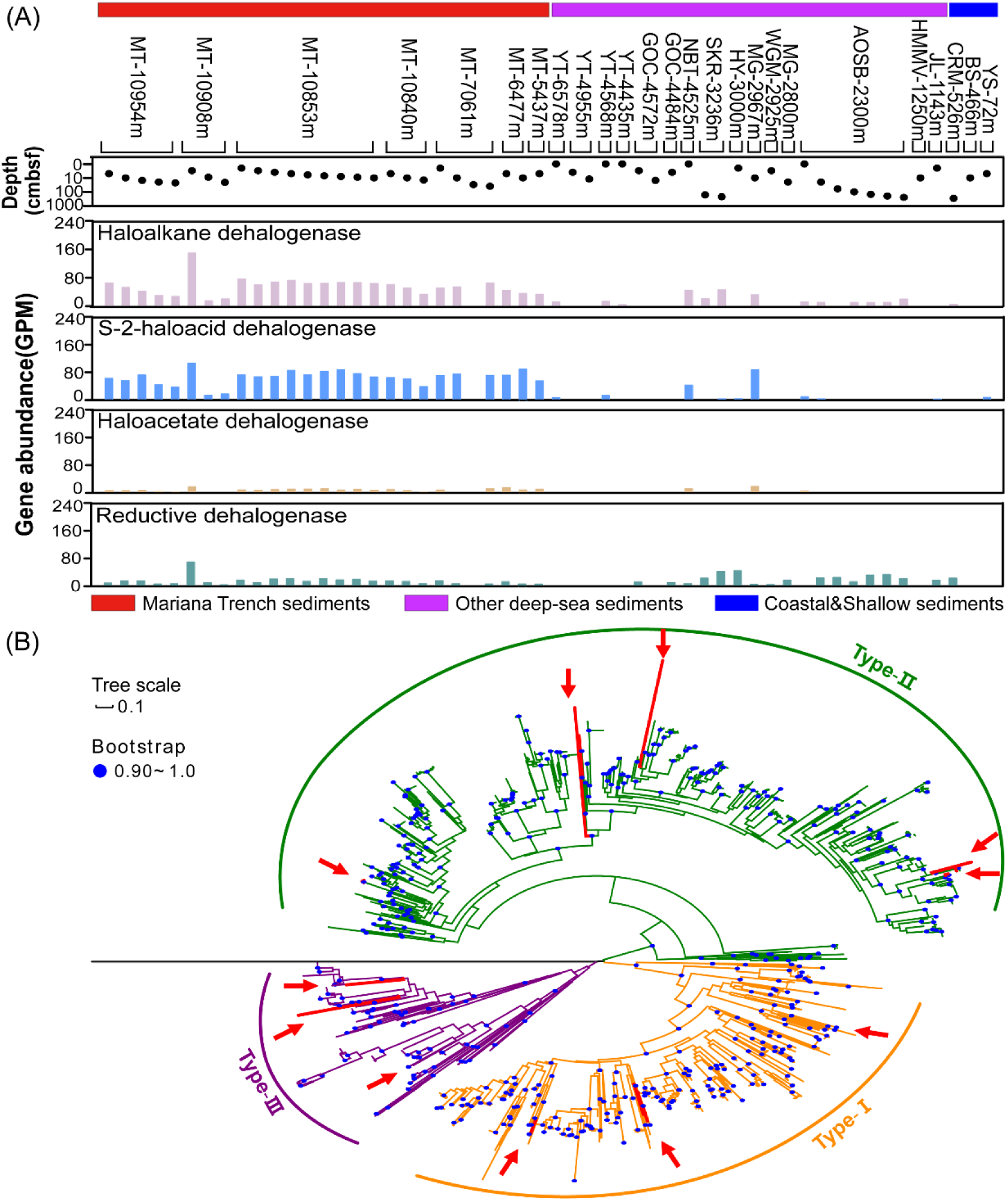
(A) Relative abundance of different types of dehalogenases in the metagenomes from the global ocean. The values were expressed as genes per million (GPM). (B) Maximum-likelihood phylogenetic tree of haloalkane dehalogenase identified from the sediment of the Mariana Trench. Red arrows indicate the location of previously characterized haloalkane dehalogenase sequences from the Uniport database. A sequence of epoxide hydrolase (accession no. P07099) was used as the root.

### Prevalence of dehalogenation metabolism among hadal trench microorganisms

In this study, a total of 2544 metagenome-assembled-genomes (MAGs) were reconstructed from the 27 metagenomes derived from the Mariana Trench sediments. After dereplication at 95% average nucleotide identity (ANI), 285 representative genomes were obtained. Functional annotation and homology confirmation revealed that over half of the trench microbial taxa, specifically 50.2% (143 MAGs), harbored genes encoding dehalogenases (Fig. 2A; Table S3). These MAGs showed contamination < 4.99%, the genome size ranged from 0.58 to 5.31 Mbp, the GC content ranged from 59.4 to 65.4%, and 79 of them had a genome completeness greater than 80% (Fig. 2A; Table S3). Among the 143 MAGs, 78 MAGs contained genes for S-2-haloacid dehalogenase, 11 for haloacetate dehalogenase, 79 for haloalkane dehalogenase, and 23 for reductive dehalogenase. While the majority of the MAGs harbored a single type of dehalogenase, 40 MAGs contained genes for multiple dehalogenase types, with one MAG from the Chloroflexota phylum (MT_10840m_10-14cm.31) possessing genes for all four dehalogenases (Fig. 2A). Phylogenomic analysis and taxonomic classification against the GTDB database revealed that the 143 MAGs with dehalogenation potentials were distributed across 16 bacterial phyla and 20 classes, including 60 novel species and 15 novel genera (Fig. 2A; Table S3). By comparing against the mibPOPdb database, which is a manually curated integrative resource for microbial bioremediation of persistent organic pollutants (POP) ^38, 39^, 11 of the 16 phyla were reported to have the capability of dehalogenation for the first time (Fig. 2A), thereby greatly enhancing the current understanding of the diversity of dehalogenation microorganisms. The MAGs encoding dehalogenases were found to be widely distributed across all analyzed MT sediment samples, with the total relative abundance of dehalogenation microorganisms constituting between 8% and 21% of the metagenome reads (Fig. 2B). Majority of the MAGs (96 MAGs) showed high occurrence frequency, being present in over 80 % of the MT sediment samples (Fig. 2B, Table S4). These findings suggest that the dehalogenation microorganisms were both prevalent and dominant in the microbial communities of the trench sediments.

Functional annotation utilizing the KEGG, COG, and PROKKA databases revealed that the dehalogenation population, represented by the 143 dehalogenation MAGs, contained gene sets that encode pathways for the degradation of various HOC compounds with distinct molecular structures. Specifically, the population was associated with the degradation of 18 compounds categorized into groups such as chlorocyclohexane and chlorobenzene, chloroalkane and chloroalkene, fluorobenzoate, as well as polychlorobiphenyl (PCB) (Fig. 3, Fig. S3, Table S5). These represent nearly all documented degradation pathways for HOCs in the KEGG database, suggesting the wide range of substrate spectra for the dehalogenation population identified in hadal trench environments. Majority of the HOCs compounds (15 of 18), e.g. γ-hexachlorocyclohexane, 2,6-dichloropenol, 2,4-dichlorophenoxyaceate, 2,4-dichloroaniline, cis-dichloropropene, trans-dichloropropene, 1,2-dichloroethane, dichloromethane, 2-fluorobenzoate, 3-fluorobenzoate, 4-chlorobenzoate, and 4-chlorobiphenyl, could be completely degraded via entering the TCA cycle (Fig. 3, Table S5). Furthermore, coding genes for each of the functional enzymes involved in these degradation pathways was identified across various genomes from multiple phyla (Fig. 3, Table S5), indicating a functional redundancy that facilitates the efficient degradation of substrates ^40^. The exact molecular composition of HOCs present in trench sediment remains undetermined. Nevertheless, as previously discussed, they are hypothesized to comprise a mixture of both anthropogenic and natural HOCs. The anthropogenic HOCs are characterized by low structural diversity ^16^, with majority of their degradation pathways being annotated in our analysis (Fig. 3, Table S5). In contrast, natural HOCs exhibit a considerably higher diversity in their molecular structures ^16, 17^. However, as the primary source of organic matter in deep-sea and hadal trenches, phytoplankton and sinking particles have been identified as containing HOCs that were predominant by aliphatic forms ^19^. Considering the diverse degradation pathways identified in the MAGs, the dehalogenation population possess the potential to efficiently utilize HOC substrates present in hadal trenches.

**Figure 3.**
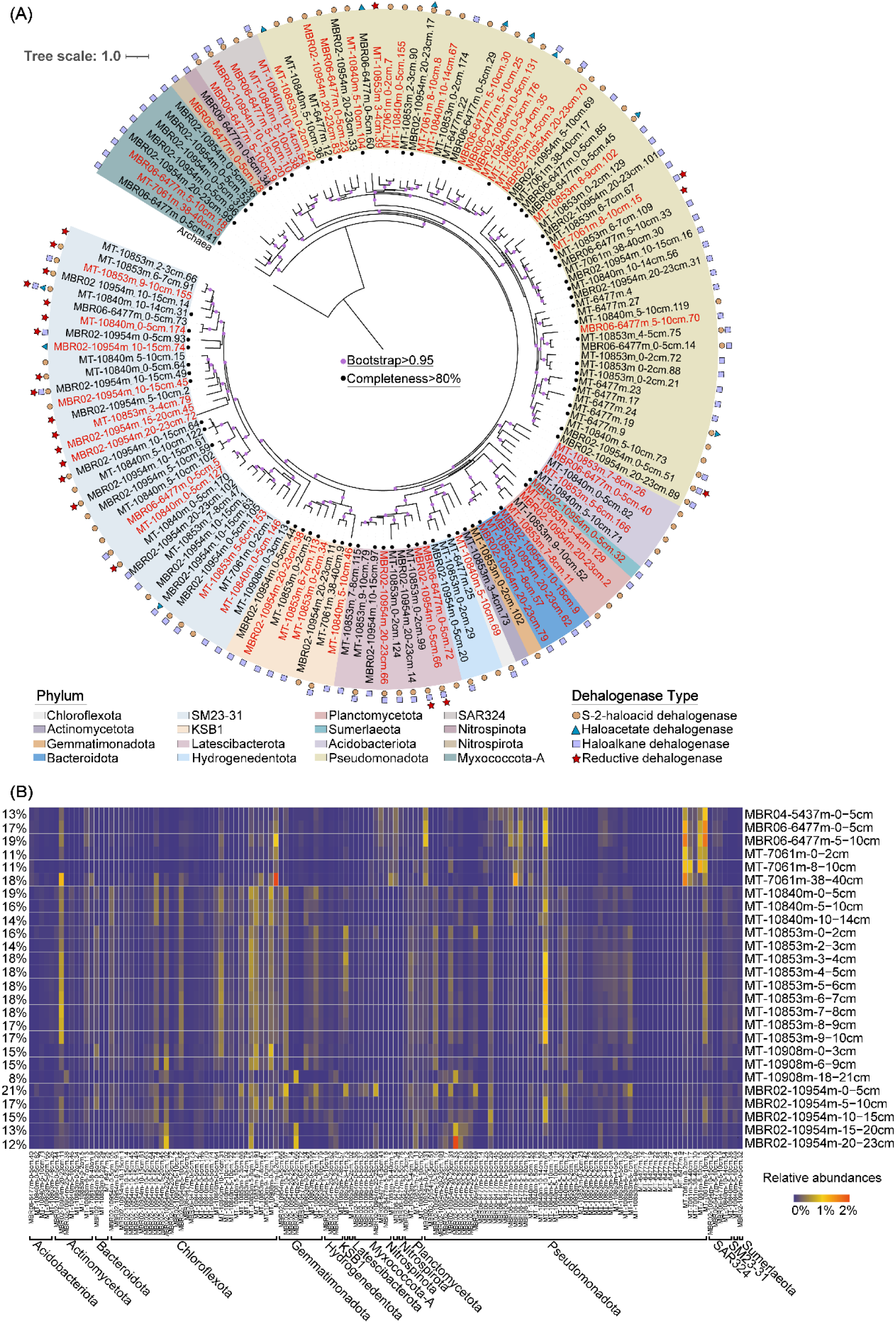
(A) Maximum-likelihood phylogenomic tree of the 143 MAGs containing genes encoding dehalogenase. An archaeal genome (accession no. GCF_004799605.1) was utilized as the root. The colored backgrounds indicate the phylum level classification, and different symbols indicate the types dehalogenase genes each MAGs contained. (B) The relative abundance of each MAG in the metagenomes from different sediment samples of the Mariana Trench. The numbers at the lefthand side indicate the total relative abundance of the 143 MAGs.

In addition to direct degradation of HOCs, the dehalogenation population showed potentials to contribute to other ecological functions. Firstly, the degradation pathways of HOCs annotated involved were generally long, with many steps and produce series of intermediates (Fig. 3). These intermediate compounds, with lowered recalcitrancy compared to their original compounds, can also be substrates for other heterotrophic taxa, supporting the potential role of the dehalogenation microorganisms as mediators of microbial interactions within hadal trenches ^16, 41^. Furthermore, the dehalogenation microbial populations also showed the capability to drive the biogeochemical cycling of multiple key elements other than halogen and carbon. Particularly, the sulfur oxidation seemed to be common in the dehalogenation population, with 49-87% of the MAGs harbored gene sets for different steps of sulfur oxidation (Fig. 3, Fig. S3, Table S5). Moreover, 43% and 45% of the MAGs also showed potentials of iron reduction and iron oxidation, respectively (Fig. 3, Fig. S3, Table S5). Overall, the findings demonstrate a significant role of dehalogenation microorganisms in the functions and structures of hadal trench ecosystems.

### Active HOCs degradation driven by hadal trench microorganisms

The degradation activity of HOCs by hadal trench microorganisms was demonstrated by inoculating typical HOCs compounds—PCB and γ-HCH—into sediments from the Mariana Trench (MBR06, 6477 m depth) and incubating them for 270 days under high pressure (60 MPa) and low temperature (4 °C) to simulate *in-situ* conditions. Time-series sampling and monitoring revealed a rapid decrease of substrate abundance in all treatments containing trench microorganisms (non-sterile systems), whereas no significant changes occurred in the control systems (sterile sediments) (Fig. 4A). For both substrates, the most substantial decrease took place within the first 30 days, leaving less than 40% of the original material. PCB levels continued to decline throughout the 270-day period, whereas γ-HCH remained relatively stable after day 30. Reads recruited to metagenomes and metatranscriptomes indicated that the dehalogenation population, i.e., the 143 MAGs with dehalogenation potential, were present and actively expressed in the microcosms (Fig. 4B). Dominant dehalogenation MAGs included those from the orders UBA1845 (p_Planctomycetota), Hydrogenedentiales (p_Hydrogenedentota), SAR324 (p_SAR 324), Longimicrobiales (p_Gemmatimonadota), UBA5794 (p_Actinomycetota), UBA9160 (p_Myxococcota), Woeseiales, Pseudomonadales, Kiloniellales (p_Proteobacteria), and SM23-28-2, UBA1151, SAR202, UBA3496, DSTF01 (p_Chloroflexota) (Fig. 4B). Among these, MAGs from the order Pseudomonadales were the most abundant in the microcosms (Fig. 4B). The majority of functional genes associated with HOC degradation pathways were found in high relative abundance in the metagenomes of the microcosms (Fig. 4C, Table S6), with over half of these genes being actively expressed in the transcriptome (Fig. 4D, Table S6). Notably, genes such as *dhaA*, *EC:1.1.1.1*, *clcD*, *catB*, *EC1.14.13.-*, and *ALDH*, which encode haloalkane dehalogenase, alcohol dehydrogenase, carboxymethylenebutenolidase, muconate cycloisomerase, 2-octaprenylphenol hydroxylase, and aldehyde dehydrogenase, respectively, exhibited high transcriptional abundance (Fig. 4D). Overall, the results from the chemical and microbial analyses in the microcosm experiments indicated rapid degradation of typical HOCs, alongside active expression of microorganisms and functional genes related with dehalogenation metabolism. This suggests that hadal trench microorganisms actively metabolize HOC under the high pressure and low temperature conditions. However, this cultivation experiment was limited to two specific human-made halogenated organic compounds. Future research is necessary to explore the metabolic activity of deep-sea microorganisms on other types of HOCs, particularly natural ones. Additionally, since this experiment utilized enriched sediment microorganisms in a laboratory setting, the degradation rates observed may differ from those in natural environments. Future in-situ cultivation studies are needed for a more accurate evaluation of the degradation rates and flux of HOCs.

**Figure 4.**
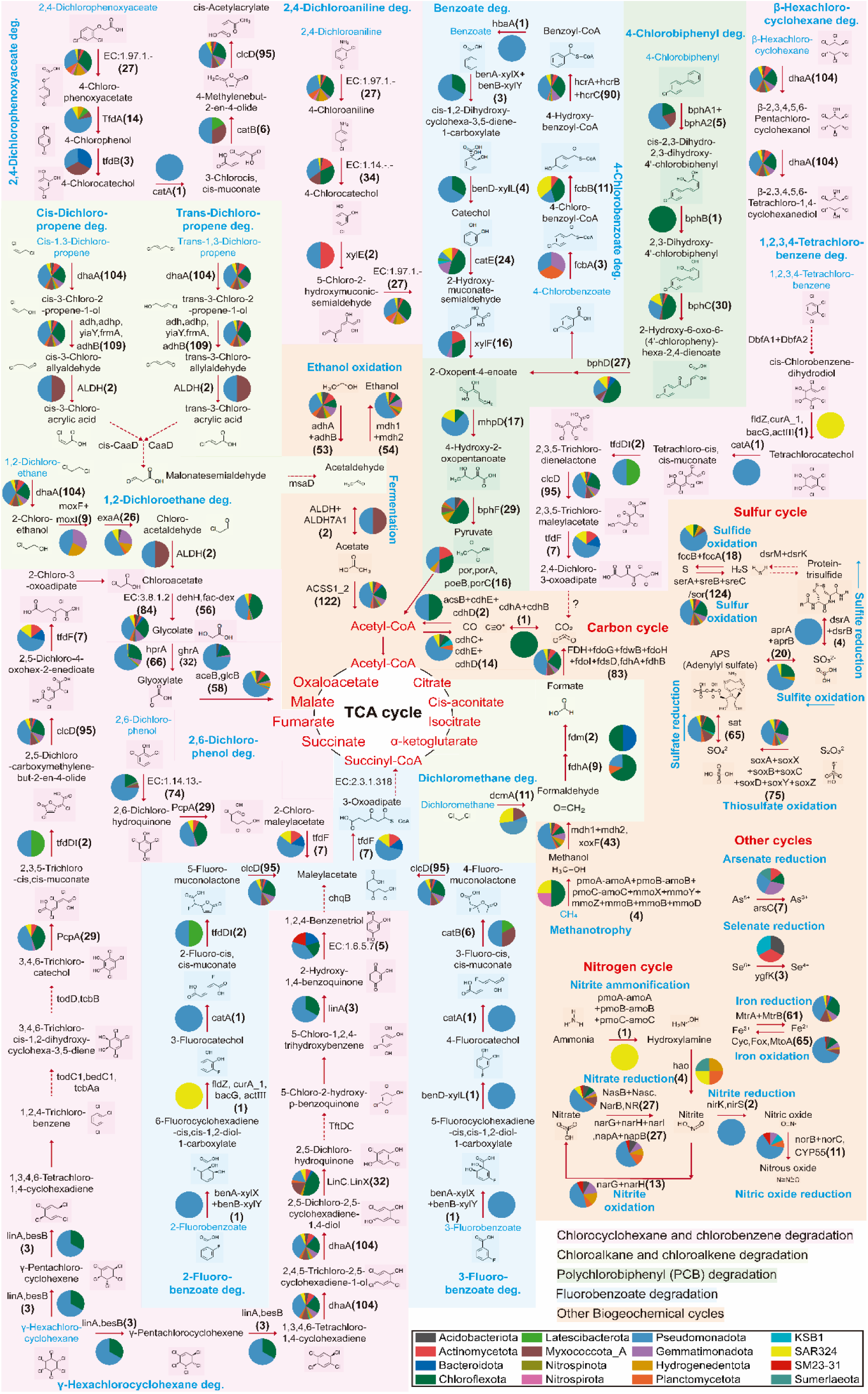
The detailed pathways for the HOCs degradation and related biogeochemical cycling by the dehalogenation microbial taxa (the 143 MAGs) identified from the sediment of the Mariana Trench. The name of the encoding gene involved in each step of degradation was shown, and the value in the brackets shows the number of MAGs containing the gene. The pie charts beside the gene names indicate the phylum level classification and proportion of the MAGs containing the gene.

**Figure 5.**
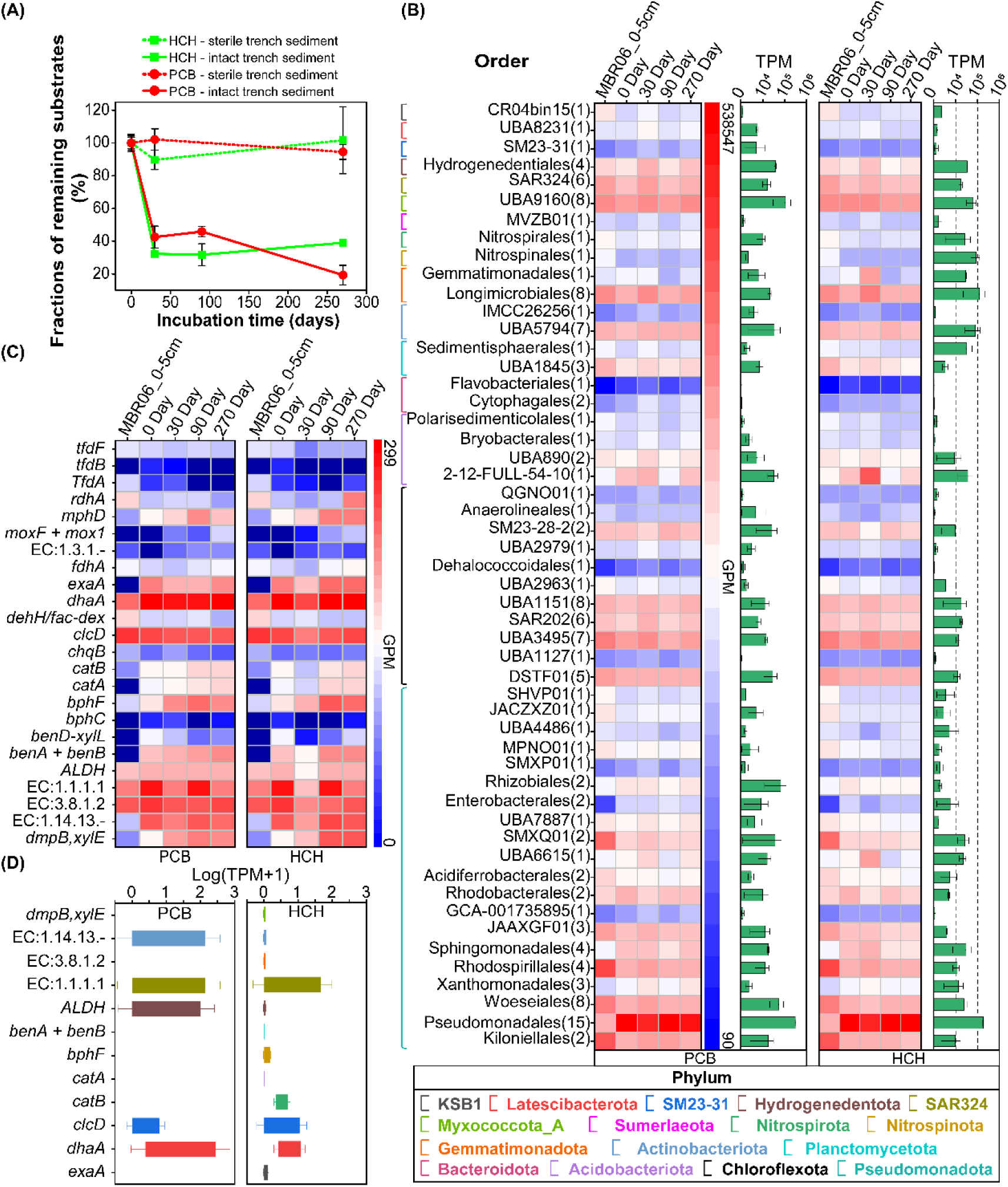
Degradation of typical HOCs compounds by microbial community from the sediment of the Mariana Trench, tested using the simulated high-pressure microcosms (A) Decreasing of both substrates with incubation time in the treatments with intact trench sediments. (B) Relative abundance (GPM) and transcription activities (TPM) of the MAGs containing dehalogenase during different times of the incubation. (C) and (D), relative abundance and transcription activities of functional genes related with pathways for degradation of HOCs.

### Implications for deep-sea biogeochemical cycling and environmental bioremediation

The deep-sea environments constitute a significant portion of the Earth’s microbial biomass and production, thereby playing a crucial role in regulation of global carbon cycling and climate change^1^. However, there remains a limited understanding of the metabolic processes of deep-sea microorganisms and the carbon turnover mechanisms they facilitate. On the other hand, over the past few decades, dehalogenation processes and the microorganisms involved have attracted a wide range of research interests, primarily due to their potential applications in the bioremediation of dioxin pollutants ^42^. Nevertheless, their ecological roles in carbon cycling of the natural environments have largely been overlooked ^16, 41^. This study presents novel evidence showing a surprisingly high concentration of HOCs within the TOC pool of deep-sea trenches. It also highlights the enrichment and prevalence of HOCs metabolic potentials across a diverse array of microbial taxa, as well as the active degradation of typical HOCs by microorganisms in the Mariana Trench under simulated *in-situ* conditions. These results indicate that HOCs may serve as important substrates for microorganisms inhabiting trench environments, and that the metabolism of HOCs may represent a common strategy for the survival of microorganisms in harsh conditions of the hadal trenches, such as the presence of recalcitrant organic matter. The findings not only enhance the current understanding of microbial processes and survival strategies in extreme marine environments, but also underscore the significance of HOCs degradation in the microbial carbon turnover in the Mariana Trench. Given that trenches are “hotspots” for microbial carbon turnover and early diagenesis in the deep ocean ^4^, the observed active degradation process of HOCs in the MT sediments is believed to have great contribution to the deep-sea organic carbon remineralization. Furthermore, our results also showed that dehalogenases present in all of the microbial metagenomes from various habitats across the global ocean (Fig. 2), with potential dehalogenation microorganisms, such as Chloroflexi, being widespread and dominant in the sediments of the hadal trenches and other deep ocean habitats ^43, 44^. Dehalogenation activities have even been recorded in deep subseafloor sediments down to 358 m below the seafloor ^45, 46^. Although research on this topic remains limited, existing evidence suggests that HOC metabolism is likely widespread in deep-sea environments. This implies that the microbial degradation of HOCs may play an important yet overlooked role in the global deep-sea carbon cycle, warranting further investigation in future studies.

Conversely, HOCs have been regarded as hazardous pollutants pose substantial environmental risks, and they are believed to ultimately become sequestered in deep-sea sediments due to their persistent nature ^15, 20^. The findings of this study demonstrate that HOCs can serve as substrates for a diverse array of microbial taxa in deep-sea environments, where they can be effectively degraded. This provides valuable new insights into the fate and environmental implications of HOCs within deep-sea ecosystems. Additionally, the high diversity of dehalogenation microorganisms along with the unique phylogenetic traits of dehalogenase identified in hadal trenches, represent promising resources for bioremediation efforts.

## Material and Methods

### Sampling sites and sample processing

Sediment samples from MBR02 (11.327°N,142.188°E, water depth 10901 m), MBR04 (10.761°N,142.274°E, water depth 5842 m), MBR05 (10.979°N, 141.950°E, water depth 6957), and MBR06 (10.813°N,141.180°E, water depth 6477 m) of the Mariana Trench were taken during the R/V *Tansuoyihao* TS15 cruise on December 2019 (Fig. 1). Samples were taken by a box corer attached to a Hadal Lander. The procedures of the sample processing were the same as in ^10, 26^: after recovering onboard, the sediment samples were immediately subsampled using sterile polypropylene corers, and most of the cores were stored under -80 °C until analysis. The pore water was collected at 1–2 cm intervals using Rhizon samplers. Aliquots for DOM analysis were stored in a pre-combusted (525 °C for 4 h) brown glass bottle, sealed with an acid-cleaned Teflon-lined cap, and frozen at −20 °C.

### Measurement of halogenated organic matter and geochemical parameters

The method for sample pretreatment and HOCs measurement followed our previous study Wei et al.,^27^. Briefly, sediment cores for HOCs measurement were thawed and divided at 2 cm interval along the vertical depth profile. The most outer layer of each depth fraction was removed to avoid halogen contamination, and the remaining sample was freeze-dried and ground. Twenty milligram sediment was then weighted out from each sample and mixed with 50 mL sodium nitrate washing solution, shaking for 2 h at 140 rpm and then stay overnight to remove inorganic halogens ^27^. The sediment slurry was then filtered through a polycarbonate (PC) membrane to collect insoluble fraction of the sediment. Eighty milligram of activated carbon was added to the filtrate and shake for 2 h at 140 rpm. The activated carbon with dissolved organic matter adsorbed was collected on a PC membrane by filtration. The two PC membranes were then subject to combustion for measurement of content of insoluble and absorbable HOCs, respectively. The protocols for combustion and detection of different halogen ions by IC were the same as Wei et al., ^27^. All labware utilized were carefully selected and pretreated according to Wei et al. (2024) to secure that they were free of halogen contamination. Three replicates were taken for each sample. Triplicate sample processing blanks and analytical blanks were involved in each batch of experiment to indicate the presence and level of the background contamination. All measured values were calibrated by subducting the background values in blank controls from the same batch of experiment.

Content of TOC (wt.%) was determined with a TOC analyzer (Multi N/C 3100 Analytik Jena) according to the procedure of Xu et al. ^47^. Carbon isotopic composition of TOC was determined according to Luo et al., ^32^, using high temperature combustion on a Vario Pyro Cube Elemental Analyzer connected to an Isoprime 100 Continuous Flow Isotope Ratio Mass Spectrometer. Stab le isotope results were reported using the per mil notation (δ, ‰) relative to the V-PDB standard. Pore-water DOC concentrations were determined by a high-temperature catalytic combustion method using a Shimadzu TOC-L total carbon analyzer with a precision of ±3% ^9^. Inorganic nutrients (NO_3_^−^, NO_2_ ^−^, NH_4_ ^+^, and PO_4_ ^3−^) were determined using a QuAAtro autoanalyzer (Seal Analytical, United Kingdom) ^48^.

### Metagenomic sequencing and public data collection

Sediment cores were thawed on ice and were depth fractioned at 5 cm intervals (2019 cruise). Total genomic DNA was extracted from 10-20 g of sediments from each depth fraction using the PowerMax^®^ Soil Kit (MoBio Laboratories, USA), and then purified by DNeasy^®^ PowerClean Pro Cleanup Kit (Qiagen, Germany). DNA fragment libraries were prepared by shearing genomic DNA from each sample and were then subjected to metagenomic sequencing on the illumine MiSeq platform in Majorbio Bio-Pharm Technology Co. Ltd (Shanghai, China). In total, 8 metagenomes were successfully sequenced and generated 335.4 Gb of raw reads. The data size for individual metagenomes ranged from 27.2 - 50.1 Gb, with an average value of 41.9 Gb. The metagenomes were combined with 46 additional deep-sequenced metagenomes from NCBI database (including 19 metagenomes from sediments of the Mariana Trench and 27 from other habitats of the global ocean) (Table S1), and were co-analyzed for distribution of dehalogenase and related microorganisms.

### Assembly, functional annotation, and dehalogenase genes identification

Metagenomic reads from each sample were quality filtered using Trimmomatic v. 0.38 ^49^ with parameters specified as “LEADING:30 TRAILING:30 CROP:90 HEADCROP:10 SLIDINGWINDOW:4:25 MINLEN:50,” and were separately assembled with metaSPAdes v. 3.15.5 (-m 700) ^50^. Gene predictions were performed against the assembled contigs using Prodigal v. 2.6.3 (-p meta) ^51^, and genes were then dereplicated to construct the non-redundant gene database using CD-HIT v. 4.8.1 with default parameters ^52^. Functional annotation was performed using BlastKOALA against the KEGG database with default parameters ^53^. The gene sequences annotated as any types of dehalogenases were extracted and confirmed using BLASTp search against UniProt protein database ^54^.Only the sequences showed ≥ 60% identity, ≥ 90% coverage and E values < 10 ^-5^ with existing dehalogenase sequences in the UniProt database were selected for downstream analysis.

### Relative abundance determination and phylogenetic analysis of dehalogenase

Clean reads from each metagenome were mapped back to confirmed dehalogenase genes sequences using BWA v. 0.7.17 ^55^. BWA output was then converted to a sorted and indexed bam file using SAMtools v. 1.9 ^56^.The number of mapped reads and length of each gene sequence from the bam file was utilized for calculation of the length normalized gene abundance, which was expressed as GPM (genes per million). The relative abundance of each type of dehalogenase was calculated by summing the GPM values of the confirmed gene sequences.

Phylogenetic analysis was conducted to reveal the differences between the hadal trench dehalogenase and the existing counterparts. A total of 2300 BLASTP confirmed haloalkane dehalogenase gens sequences were extracted from the 28 Mariana Trench metagenomes, translated in to protein sequences, and co-analyzed with 37 previously characterized haloalkane dehalogenase sequences from the Uniport database. The sequences were aligned with MAFFT v. 7.467 ^57^ and trimmed with trimAl software ^58^, with default parameters. A maximum-likelihood phylogenetic tree was constructed with IQ-Tree ^59^with 100 bootstrap replicates.

### Genome binning and dereplication

The quality-filtered reads were mapped back to the assembled contigs using Bowtie2 (v. 2.3.4.1) ^60^, and coverage was determined according to the mapping results with the jgi_summa-rize_bam_contig_depths script ^60^. Metagenome binning was conducted for assemblies longer than 2500 bp using MetaBAT v. 2.12.1 ^61^ with default parameters. Quality of the MAGs was assessed by CheckM v. 1.1.2 using the line-age_wf workflow ^62^, and only MAGs with completeness > 50% and contamination < 5% were kept for further analysis. Redundant bins were subsequently dereplicated using dRep v. 2.3.2 ^63^ at 95% average nucleotide identity (ANI) (all other parameters were set to the default), and MAG with the highest quality was selected from each cluster for downstream analysis. The genome size was estimated by dividing the size of the MAG by its estimated completeness.

### Taxonomic assignment, identification of dehalogenation taxa and calculation of their relative abundance

Taxonomic classification of the MAGs was determined using GTDB-Tk, which is based on the phylogenetically calibrated Genome Taxonomy Database (GTDB) ^64^. Coding sequences in the MAGs were predicted using Prodigal v. 2.6.3 with default setting ^51^. Functional annotation was performed by using BlastKOALA against the KEGG database with default parameters ^53^. The annotation results were manually examined to screen for the MAGs with dehalogenase annotated. The sequences of dehalogenase genes were then extracted from each candidate MAG and confirmed using BLASTp search against UniProt protein database. Only the MAGs containing gene sequences showing ≥60% identity and ≥ 90% coverage with existing dehalogenase in the UniProt database were selected as the potential dehalogenation microorganisms.

Phylogenomic tree was constructed for the confirmed dehalogenation MAGs, using the 43 universal single-copy genes (SCGs) used by CheckM ^62^. Protein sequences of the SCGs were identified using HMMER v. 3.1b2 ^65^ with default parameters, individually aligned with MAFFT v. 7.467 ^57^ and then concatenated. Phylogenomic tree was constructed based on the alignment using FastTree2 v. 2.1.11 ^66^, with a JTT model, a gamma approximation and 100 bootstrap replicates. Archaeal genome (GCF_004799605.1) was utilized as the root. The phylogenomic tree was visualized with iTOL ^67^.

Relative abundances of the dehalogenation MAGs were calculated by mapping the trimmed reads to the manually curated and refined MAGs with CoverM (v.0.6.1; https://github.com/wwood/CoverM) in genome mode including the dereplication flag using the default aligner Minimap2 (v.2.21; https://docs.csc.fi/apps/minimap2/) in short-read mode, discarding unmapped reads. The final relative abundance of each MAG is the percentage of the MAG in the mapped fraction of each sample.

### Reconstruction of metabolic network for dehalogenation microorganisms

Systematic metabolic capabilities of the dehalogenation microbial taxa were explored by systematic annotation of MAGs containing dehalogenase genes. Coding sequences in the MAGs were annotated against the KEGG ^53^, Cluster of Orthologous Groups (COG) ^68^, as well as PROKKA databases ^69^, following the steps in Liu et al. ^10^. The pathways related with energy metabolism and biogeochemical cycling were also annotated with METABOLIC ^70^. Metabolic network was reconstructed for each of the MAGs by summarizing the annotated pathways.

Coding sequences in the MAGs were annotated against the KEGG^53^, Cluster of Orthologous Groups (COG)^68^, as well as PROKKA databases^69^, following the steps in Liu et al.^10^. The pathways related with energy metabolism and biogeochemical cycling were further annotated with METABOLIC^70^. Metabolic network was reconstructed for each of the MAGs by summarizing the annotated pathways.

## High-pressure microcosms

### Microcosm set up and sampling

Sediments were taken from site MBR06 (water depth 6477 m) of the Mariana Trench using a box corer during RV Tansuoyihao TS15 cruise on December 2019. After removing the outer layers, around 2.0 kg sediment were taken by pre-combusted (450 °C, 4 h) stainless steel scoops. Sediment samples were transferred into sterile incubation bags (polypropylene) and stored under 4 °C for 20 months, which leads to the enrichment of dehalogenation microbial taxa (relative abundance of dehalogenation microbial taxa increased from < 20% to 53% of total mapped metagenomic reads).

After thorough mixing with a pre-combusted stainless steel scoop, the sediments were divided into polypropylene incubation bags and mixed with 0.2 µm filtered seawater taken from 6000m depth of the Mariana Trench. Each of the 50 ml incubation bag contained 15 g sediment and 30 ml filtration sterilized seawater, and then either 1.5 mg of 4-Chlorobiphenyl (4-PCB) or 3 mg of γ-Hexachlorocyclohexane (γ-HCH) was amended as the representatives of anthropogenic HOCs with low or high numbers of halogen atoms, respectively. No-substrate controls and no-microorganism controls were set up simultaneously with the experimental groups. No-substrate controls are incubation bags contained only sediment and sterilized seawater, and without halogenated organic matter substrates. No microorganism controls are incubation bags containing autoclaving sterilized sediments and filtration sterilized seawater, and either type of halogenated organic substrates with the same ratios as the experiment groups. All of the incubation bags for experimental groups and the controls were put into high pressure incubation vessels, and were incubated under 60 MPa and 4 °C to simulate the *in-situ* pressure and temperature. Triplicate incubation bags were taken for each treatment at day 0, day 30, day 60, day 90, and day 270 to monitoring the changes of organic substrates and microbial metabolisms.

### 4-PCB and γ-HCH measurement

Around 15 mL of sediment slurry from each incubation bags were freeze dried and ground. After mixing with 100 µL surrogate standard solution (2,4,5,6-Tetrachlorom-xylene, 500 mg/L), the sample was transferred into Accelerated Solvent Extraction (ASE) cells for extraction with (1:1, v/v) (72 °C, 20 h for 4-PCB, and 65 °C, 22h for HCH). The yielded extract from each cell was then evaporated to a final volume of 1 mL and then cleaned up in CNWBOND Florisil PR SPE cartridge (60-100 mesh, 1 g/6 mL). First, 1 g of Na_2_SO_4_ was added to the cartridge and 10 mL of n-hexane was added to pre-condition the column. Then the extract was transferred into the column and eluted by 10 mL of acetone:n-hexane (1:1). The collected effluent was concentrated to 5 mL. Before instrumental analysis, 100 µL of the effluent was transferred into amber GC vials, and 50 µL of injection standard (50 mg /L, 1-Bromo-2-nitrobenzene for 4-PCB analysis, Pentachloronitrobenzene for HCH analysis) was added to calculate recoveries.

The 4-PCB and HCH were analyzed by using gas chromatograph (Agilent 7890B; DB-5 column, 15m× 0.32mm×0.25 μm) equipped with 63Ni-ECD detector (for analysis of HCH) and MSD detector (for analysis of PCB). Samples (1 μL) were injected in the splitless injection mode with high purity nitrogen as carrier gas (1 mL/min for 4-PCB analysis, and 2 mL/min for HCH analysis). For analysis of 4PCB, the temperature of the column was set at 40°C for 2 min, then raised to 280°C (20°C·min^−1^), and finally retained at 280°C for 5 min. The injection port and detector temperatures were kept at 270°C and 280°C, respectively. For analysis of HCH, the temperature of the column was set at 100°C for 2 min, raised to 220°C (15°C·min^−1^) and retained for 5 min, then raised to 260 °C (15°C·min^−1^) and retained for 20 min. The injection port and detector temperatures were kept at 220°C and 300°C, respectively.

The quality control steps included sample processing blanks, limit of detection, recovery tests, replicates, and standard curves. Sample processing blanks were involved for each batch of experiment to indicate possible contamination. All blanks were below the limits of detection (LOD). The LODs were set to be triple the standard deviation of the blanks, which were averagely 0.1 µg/g for PCB and 0.01µg/g for HCH. Spiked recovery tests of the standards showed acceptable recoveries ranging from 81.80% to 85.40% for 4-PCB, with the relative standard deviation (RSD) less than ±2.57%, and from 91.59% to 94.63 % for HCH, with the RSD less than ± 4.74% . The standard curves had high degree of linearity (R^2^ > 0.999) for all the compounds.

### Metagenomic and metatranscriptomic analysis

Total DNA and RNA were coextracted from each of the microcosm samples using PowerSoil Total RNA Isolation Kit and DNA Elution Accessory Kit (MoBio Lab, United States). The concentration and quality of DNA and RNA were determined using both Qubit 4.0 (Thermo Fisher Scientific, Waltham, USA) and Nanodrop One (Thermo Fisher Scientific, Waltham, USA). Quality of RNAs were also determined through the Agilent 4200 system (Agilent Technologies, Waldbronn, Germany). All of the DNA samples were directly subject to library preparation using the ALFA-SEQ DNA library Prep Kit (Findrop Biosafety Tech., China). The RNA samples went through whole transcriptome amplification using RNA REPLI-g Cell WGA & WTA Kit (Qiagen, Germany), and subject to library construction using ALFA-SEQ RNA library Prep Kit II (Findrop Biosafety Tech., China). The DNA libraries and RNA libraries were then subject to metagenome and metatranscriptome sequencing, respectively, in Magigene Biotechnology Co. Ltd (Guangzhou, China) using Illumina novaseq6000 (PE150 sequencing).

The sequencing was successful for all of the metagenomes. Metagenomic reads from each sample were quality filtered, assembled, and genes were predicted and dereplicated following the same procedure as mentioned earlier. Functional annotation was performed against the KEGG, COG and PROKKA databases. The gene sequences involved in any of the HOCs degradation pathways were extracted and confirmed via BLASTp. Clean reads from each metagenome were mapped back to predicted genes using BWA v. 0.7.17 and the gene length normalized relative abundance of each gene were expressed as GPM. The relative abundances of the dehalogenation genomes were estimated by mapping the metagenomic reads to sequences of the corresponding MAGs and expressed as GPM.

Duplicated metatranscriptomes were successfully sequenced for both PCB and HCH microcosm samples taken from 90^th^ day. Metatranscriptomic raw reads were quality filtered using Trimmomatic v. 0.38 ^49^, and non-coding RNAs were removed using sortmerna v. 4.2.0 ^71^. Clean reads of each metatranscriptome were then mapped to the predicted protein coding genes of the metagenome from the same sample. The transcription level of each gene was expressed as transcripts per million (TPM). Similarly, the transcription level of the dehalogenation genomes was estimated by mapping the metatranscriptome clean reads to sequences of the corresponding MAGs and expressed as TPM.

## Acknowledgements

This work was supported by the National Natural Science Foundation of China (grant numbers 42276149, 92251303) and the Shanghai Frontiers Research Fund of the Hadal Biosphere. We also acknowledge organizers, technological teams and crews in the cruises of M.V. Zhangjian and R.V. Tansuoyihao.

## References

1. Herndl, G. J., Bayer, B., Baltar, F., Reinthaler, T. Prokaryotic life in the deep ocean’s water column. Annu. Rev. Mar. Sci. 15, 461–83 (2023).

2. Du M, et al. Geology, environment, and life in the deepest part of the world’s oceans. Innovation (New York, NY) 2, 100109 (2021).

3. Jamieson A. The Hadal Zone: Life in the Deepest Oceans (2015).

4. Glud RN, et al. Hadal trenches are dynamic hotspots for early diagenesis in the deep sea. Communications Earth & Environment 2, 21 (2021).

5. Nunoura T, et al. Hadal biosphere: Insight into the microbial ecosystem in the deepest ocean on Earth. Proceedings of the National Academy of Sciences of the United States of America 112, E1230–E1236 (2015).

6. Zhou YL, Mara P, Cui GJ, Edgcomb VP, Wang Y. Microbiomes in the Challenger Deep slope and bottom-axis sediments. Nature Communications 13, 1515 (2022).

7. Thamdrup B, et al. Anammox bacteria drive fixed nitrogen loss in hadal trench sediments. Proceedings of the National Academy of Sciences of the United States of America 118, 46 (2021).

8. Chen P, et al. Revealing the full biosphere structure and versatile metabolic functions in the deepest ocean sediment of the Challenger Deep. Genome Biology 22, 207 (2021).

9. Hu TC, et al. Probing sedimentary DOM in the deepest sector of Earth’s surface. Marine Chemistry 237,104033 (2021).

10. Liu RL, et al. Novel Chloroflexi genomes from the deepest ocean reveal metabolic strategies for the adaptation to deep-sea habitats. Microbiome 10, 1–17 (2022).

11. Guan H, et al. Composition and origin of lipid biomarkers in the surface sediments from the southern Challenger Deep, Mariana Trench. Geoscience Frontiers 10, 351–360 (2019).

12. Liu JW, et al. Proliferation of hydrocarbon-degrading microbes at the bottom of the Mariana Trench. Microbiome 7, 47 (2019).

13. Chu MF, et al. Earthquake-enhanced dissolved carbon cycles in ultra-deep ocean sediments. Nature Communications 14, 5427 (2023).

14. Jiang HC, et al. Three-layer circulation in the world deepest hadal trench. Nature Communications 15, 8949 (2024).

15. Sanganyado E, Chingono KE, Gwenzi W, Chaukura N, Liu WH. Organic pollutants in deep sea: Occurrence, fate, and ecological implications. Water Research 205, 117658 (2021).

16. Atashgahi S, Häggblom MM, Smidt H. Organohalide respiration in pristine environments: implications for the natural halogen cycle. Environmental Microbiology 20, 934–948 (2018).

17. Gribble GW. Naturally Occurring Organohalogen Compounds-A Comprehensive Review. Progress in the chemistry of organic natural products 121, 1–546 (2023).

18. Liu YH, Wang L, Liu RL, Fang JS. Biogeochemical cycling of halogenated organic compounds in the ocean: Current progress and future directions. Deep-Sea Research Part I-Oceanographic Research Papers 205, 104237 (2024).

19. Leri AC, et al. A marine sink for chlorine in natural organic matter. Nature Geoscience 8, 620–624 (2015).

20. Wagner CC, et al. A Global 3-D Ocean Model for PCBs: Benchmark Compounds for Understanding the Impacts of Global Change on Neutral Persistent Organic Pollutants. 33, 469–481 (2019).

21. Peng GY, Bellerby R, Zhang F, Sun XR, Li DJ. The ocean’s ultimate trashcan: Hadal trenches as major depositories for plastic pollution. Water Research 168, 115121 (2020).

22. Sobek A, et al. Organic matter degradation causes enrichment of organic pollutants in hadal sediments. Nature Communications 14, 2012 (2023).

23. Cui JT, et al. Occurrence of Halogenated Organic Pollutants in Hadal Trenches of the Western Pacific Ocean. Environmental Science & Technology 54, 15821–15828 (2020).

24. Dasgupta S, et al. Toxic anthropogenic pollutants reach the deepest ocean on Earth. Geochemical Perspectives Letters 7, 22–26 (2018).

25. Jamieson AJ, Malkocs T, Piertney SB, Fujii T, Zhang ZL. Bioaccumulation of persistent organic pollutants in the deepest ocean fauna. Nature Ecology & Evolution 1, 0051 (2017).

26. Liu RL, et al. Bulk and Active Sediment Prokaryotic Communities in the Mariana and Mussau Trenches. Frontiers in Microbiology 11, 1521 (2020).

27. Wei H, Liu YH, Huang LT, Wang L, Fang JS, Liu RL. Determining the abundance, composition and spatial distribution of organohalogens in marine sediments using combustion-ion chromatography. Marine Environmental Research 199,106626 (2024).

28. Xiao WJ, Xu YP, Canfield DE, Wenzhöfer F, Zhang CL, Glud RN. Strong linkage between benthic oxygen uptake and bacterial tetraether lipids in deep-sea trench regions. Nature Communications 15,3439 (2024).

29. Zhang X, et al. The hadal zone is an important and heterogeneous sink of black carbon in the ocean. Communications Earth & Environment 3, 25 (2022).

30. Blum JD, Drazen JC, Johnson MW, Popp BN, Motta LC, Jamieson AJ. Mercury isotopes identify near-surface marine mercury in deep-sea trench biota. Proceedings of the National Academy of Sciences of the United States of America 117, 29292–29298 (2020).

31. Xie JQ, et al. First evidence and potential sources of novel brominated flame retardants and BDE 209 in the deepest ocean. Journal of Hazardous Materials 448,130974 (2023).

32. Luo M, et al. Benthic Carbon Mineralization in Hadal Trenches: Insights From In Situ Determination of Benthic Oxygen Consumption. Geophysical Research Letters 45, 2752–2760 (2018).

33. Xiao WJ, Wang YS, Liu YS, Zhang X, Shi LL, Xu YP. Predominance of hexamethylated 6-methyl branched glycerol dialkyl glycerol tetraethers in the Mariana Trench: source and environmental implication. Biogeosciences 17, 2135–2148 (2020).

34. Fang H, et al. Metagenomic analysis reveals potential biodegradation pathways of persistent pesticides in freshwater and marine sediments. Science of the Total Environment 470, 983–992 (2014).

35. Thomas AW, Lewington J, Hope S, Topping AW, Weightman AJ, Slater JH. Environmentally directed mutations in the dehalogenase system of Pseudomonas putida strain PP3. Archives of Microbiology 158, 176–182 (1992).

36. Zhao Y, Ma J, Zhang L, Tian K, Yao D, Yun Liu. Biodegradation mechanism of organochlorine pesticides: a review. Environ. Prot. Chem. Ind. 41, 551–558 (2021).

37. Koudelakova T, et al. Substrate specificity of haloalkane dehalogenases. The Biochemical journal 435, 345–354 (2011).

38. Ngara TR, Zeng PJ, Zhang HJ. mibPOPdb: An online database for microbial biodegradation of persistent organic pollutants. Imeta 1, e45 (2022).

39. Ruan Z, Xu X, Chen K, Qiao W, Jiang J. Recent advances in microbial catabolism of persistent organic pollutants. 60, 2763–2784 (2020).

40. Li DD, et al. Quantifying functional redundancy in polysaccharide-degrading prokaryotic communities. Microbiome 12, 120 (2024).

41. Yang Y, et al. Roles of Organohalide-Respiring Dehalococcoidia in Carbon Cycling. Msystems 5, e00757–e00719 (2020).

42. Adrian L, Löffler F. Organohalide-Respiring Bacteria (2016).

43. Hoshino T, et al. Global diversity of microbial communities in marine sediment. 117, 27587–27597 (2020).

44. Wu JX, et al. Biogeographic distribution, ecotype partitioning and controlling factors of Chloroflexi in the sediments of six hadal trenches of the Paciflc Ocean. Science of the Total Environment 880, 163323 (2023).

45. Futagami T, Morono Y, Terada T, Kaksonen AH, Inagaki F. Dehalogenation Activities and Distribution of Reductive Dehalogenase Homologous Genes in Marine Subsurface Sediments. Applied and Environmental Microbiology 75, 6905–6909 (2009).

46. Futagami T, Morono Y, Terada T, Kaksonen AH, Inagaki F. Distribution of dehalogenation activity in subseafloor sediments of the Nankai Trough subduction zone. Philosophical Transactions of the Royal Society B-Biological Sciences 368, 368 (2013).

47. Xu YP, et al. Distribution, Source, and Burial of Sedimentary Organic Carbon in Kermadec and Atacama Trenches. Journal of Geophysical Research-Biogeosciences 126, e2020JG006189 (2021).

48. Liu R, et al. Depth-Resolved Distribution of Particle-Attached and Free-Living Bacterial Communities in the Water Column of the New Britain Trench. Front Microbiol 9, 625 (2018).

49. Bolger AM, Lohse M, Usadel B. Trimmomatic: a flexible trimmer for Illumina sequence data. Bioinformatics 30, 2114–2120 (2014).

50. Nurk S, Meleshko D, Korobeynikov A, Pevzner PA. metaSPAdes: a new versatile metagenomic assembler. Genome Research 27, 824–834 (2017).

51. Hyatt D, Chen GL, LoCascio PF, Land ML, Larimer FW, Hauser LJ. Prodigal: prokaryotic gene recognition and translation initiation site identification. Bmc Bioinformatics 11, 119 (2010).

52. Li WZ, Godzik A. Cd-hit: a fast program for clustering and comparing large sets of protein or nucleotide sequences. Bioinformatics 22, 1658–1659 (2006).

53. Kanehisa M, Sato Y, Morishima K. BlastKOALA and GhostKOALA: KEGG Tools for Functional Characterization of Genome and Metagenome Sequences. Journal of Molecular Biology 428, 726–731 (2016).

54. Liu WG, Schmidt B, Müller-Wittig W. CUDA-BLASTP: Accelerating BLASTP on CUDA-Enabled Graphics Hardware. Ieee-Acm Transactions on Computational Biology and Bioinformatics 8, 1678–1684 (2011).

55. Li H, Durbin R. Fast and accurate short read alignment with Burrows-Wheeler transform. Bioinformatics 25, 1754–1760 (2009).

56. Etherington GJ, Ramirez-Gonzalez RH, MacLean D. bio-samtools 2: a package for analysis and visualization of sequence and alignment data with SAMtools in Ruby. Bioinformatics 31, 2565–2567 (2015).

57. Katoh K, Standley DM. MAFFT Multiple Sequence Alignment Software Version 7: Improvements in Performance and Usability. Molecular Biology and Evolution 30, 772–780 (2013).

58. Capella-Gutiérrez S, Silla-Martínez JM, Gabaldón T. trimAl: a tool for automated alignment trimming in large-scale phylogenetic analyses. Bioinformatics 25, 1972–1973 (2009).

59. Nguyen LT, Schmidt HA, von Haeseler A, Minh BQ. IQ-TREE: A Fast and Effective Stochastic Algorithm for Estimating Maximum-Likelihood Phylogenies. Molecular Biology and Evolution 32, 268–274 (2015).

60. Langmead B, Salzberg SL. Fast gapped-read alignment with Bowtie 2. Nature Methods 9, 357–U354 (2012).

61. Kang DD, Froula J, Egan R, Wang Z. MetaBAT, an efficient tool for accurately reconstructing single genomes from complex microbial communities. PeerJ 3, e1165 (2015).

62. Parks DH, Imelfort M, Skennerton CT, Hugenholtz P, Tyson GW. CheckM: assessing the quality of microbial genomes recovered from isolates, single cells, and metagenomes. Genome Research 25, 1043–1055 (2015).

63. Olm MR, Brown CT, Brooks B, Banfield JF. dRep: a tool for fast and accurate genomic comparisons that enables improved genome recovery from metagenomes through de-replication. Isme Journal 11, 2864–2868 (2017).

64. Chaumeil PA, Mussig AJ, Hugenholtz P, Parks DH. GTDB-Tk: a toolkit to classify genomes with the Genome Taxonomy Database. Bioinformatics 36, 1925–1927 (2020).

65. Eddy SR. A new generation of homology search tools based on probabilistic inference. Genome informatics International Conference on Genome Informatics 23, 205–211 (2009).

66. Price MN, Dehal PS, Arkin AP. FastTree 2--approximately maximum-likelihood trees for large alignments. PLoS One 5, e9490 (2010).

67. Letunic I, Bork P. Interactive tree of life (iTOL) v3: an online tool for the display and annotation of phylogenetic and other trees. Nucleic Acids Research 44, W242–W245 (2016).

68. Tatusov RL, et al. The COG database: an updated version includes eukaryotes. Bmc Bioinformatics 4, 41 (2003).

69. Seemann T. Prokka: rapid prokaryotic genome annotation. Bioinformatics 30, 2068–2069 (2014).

70. Zhou ZC, et al. METABOLIC: high-throughput profiling of microbial genomes for functional traits, metabolism, biogeochemistry, and community-scale functional networks. Microbiome 10, 33 (2022).

71. Kopylova E, Noé L, Touzet H. SortMeRNA: fast and accurate filtering of ribosomal RNAs in metatranscriptomic data. Bioinformatics 28, 3211–3217 (2012).

